# Benchmarking of Human Read Removal Strategies for Viral and Microbial Metagenomics

**DOI:** 10.1101/2025.03.21.644587

**Authors:** Matthew Forbes, Duncan Y. K. Ng, Róisín M. Boggan, Andrea Frick-Kretschmer, Jillian Durham, Oliver Lorenz, Bruhad Dave, Florent Lassalle, Carol Scott, Josef Wagner, Adrianne Lignes, Fernanda Noaves, David K. Jackson, Kevin Howe, Ewan M. Harrison

## Abstract

Human reads are a key contaminant in microbial metagenomics and enrichment-based studies, requiring removal for computational efficiency, biological analysis, and privacy protection. Various *in silico* methods exist, but their effectiveness depends on the parameters and reference genomes used. Here, we assess different methods, including the impact of the updated T2T-CHM13 human genome versus GRCh38. Using a synthetic dataset of viral and human reads, we evaluated performance metrics for multiple approaches. We found that the usage of high-sensitivity configuration of Bowtie2 with the T2T-CHM13 reference assembly significantly improves human read removal with minimal loss of specificity, albeit at higher computational cost compared to other methods investigated. Applying this approach to a publicly available microbiome dataset, we effectively removed sex-determining SNPs with little impact on microbial assembly. Our results suggest that our high-sensitivity Bowtie2 approach with the T2T-CHM13 is the best method tested to minimise identifiability risks from residual human reads.

## Introduction

When performing next-generation sequencing directly on human-derived microbial or viral (referred to henceforth as pathogen) samples (e.g. clinical specimens), a significant proportion of reads will originate from the human host. This problem is particularly acute in shotgun metagenomic studies when there is no or only limited depletion of host DNA. The overall proportions expected will vary depending on the protocol employed. For instance, in microbiome studies, gut microbiome studies with samples derived from faeces will typically have a low proportion of human reads (<10%), whereas saliva, nasal, skin and vaginal samples can often be majority human in composition (>90% reads) (1).

This can have two important effects, relating to both downstream analyses and legal and ethical concerns. The human genome is around a thousand-fold larger than the average bacterial genome and around 100,000-fold larger than a respiratory virus genome such as SARS-CoV-2 (∼30Kb for SARS-CoV-2 *cf.* ∼3.2Gb for the human genome). This means that a sample containing an equal balance of human and pathogen genetic material would contain more than 99.9% of human DNA (2) which can have detrimental effects on the quality of variant calls (3). Additionally, the increasing amount of human whole genome sequences or genotyping data deposited in public archives raises a risk for potential identification of the individual from whom the sample was derived. This raises numerous concerns around privacy, lack of informed consent, legal and regulatory compliance (4). This risk can present a barrier to open and timely data sharing. Removal prior to analysis is important as it ensures reproducibility with data later submitted to public archives while optimising data to mitigate erroneous calling, with the additional benefit of reducing the size of data files used for complex downstream analysis, thus reducing the energy requirement for the bioinformatics analysis (5).

Some methods seek to remove human reads *in vitro.* One method is to employ chemical depletion of host DNA. However, this can lead to issues such as incomplete removal of host DNA, loss of pathogen DNA, bias towards certain pathogen populations and increased protocol complexity, time and cost (1). Other methods enrich the microbial component, for example, 16S ribosomal (r)RNA sequencing and SARS-CoV-2 ARTIC amplicon sequencing (6); these methods will reduce the numbers of human reads to negligible levels via specific amplification of target gene(s) or genomes; however, this comes at the expense of strain-specific information in other parts of microbial genomes and can only be employed to investigate a limited number of different species.

Another method is bait capture, where a synthetic oligonucleotide probe with complementarity to target sequences is used to selectively enrich the sequencing library with particular target genes or genomes. Baits can be designed for a range of targets, including respiratory and vector-borne viruses and bacteria. This can be very useful for enhancing the signal for a group of pathogens from the background and avoids issues associated with amplification, such as primer artefacts and dropouts, however, a background of human reads is is still generated. As such, it is important to remove human reads *in silico*.

Computational Human Read Removal (HRR) methods can often be broadly divided into the following classes: alignment and classifier-based. The alignment-based methodologies involve aligning reads to a human reference and retaining only the reads which do not align. Classification-based methods involve predicting the taxonomic identity of individual reads and removing reads which are classified as human.

Various human read removal methods have previously been assessed by Bush et al. (2), who investigated several alignment and classifier-based methodologies on a benchmark of multiple simulated mixtures of bacterial species. The authors found that alignment-based methodologies more accurately located human reads compared to classification-based methods because aligners directly compare the reads with the human genome sequence. They note a trade-off in terms of computational efficiency, with alignment-based methods working on substantially longer time-scales as well as requiring greater computational resources. For short reads (<150bp) Bowtie2 followed by SNAP alignment was the most effective human read removal combination, however, they note the higher computational cost of combining the two alignment-based methods. Though the authors discuss a comparison of methodologies, they broadly implemented default parameters across the tested methodologies, however, aligners such as Bowtie2 can be further parameterised and run in alternative ways to enhance their efficacy for HRR, offering potential improvements for this use case.

Bush et al. also compared human read removal performance with aligners and classifiers against the GRCh37 (7) and GRCh38 (8) references and found no difference between methods which use the different reference genomes (2). However, since the publishing of this paper, a more complete Telomere-to-Telomere (T2T-CHM13) reference has been published, comprising the first full-length human genome sequence (9). Compared to the GRCh37 assembly, GRCh38 was an improvement on the previous best reference, but T2T-CHM13 represents a paradigm shift. The use of long-read technologies meant that the T2T-CHM13 assembly was able to resolve previously “dark” regions in the previous assembly, including centromeres, telomeres and ribosomal DNA (rDNA) arrays (9). This offers potential for large improvements in the ability of alignment-based HRR tools to improve their ability to detect human reads that previously fell into these “dark zones” and ensure their removal from the dataset.

In this study, we investigated a number of HRR methods and their capacity to separate human reads from microbial (viral and bacterial). We created a synthetic titration benchmark dataset based on a gradient of bait capture derived reads mixed with reads derived from human samples derived from the 1000 genomes project (10) across multiple populations to ensure a range of demographic representation. A read-fate analysis was performed on multiple HRR methods using the synthetic titration dataset, comprising combinations of alignment and classification based methods to identify the best performing method and investigate performance differentials. We then applied our chosen method to a publicly available microbiome data set and showed the effect that application has on the identifiability of individuals, and confirm no adverse effects of implementing this methodology on either metagenomic assembled genomes (MAG) or consensus FASTA generation.

## Methods

### Human Read Removal Modules

To identify the best practice for human read removal from microbial metagenomics data (shotgun and bait capture), we investigated the performance of multiple Human Read Removal (HRR) approaches (Table 1). These methods were selected as a representative set covering popular alignment and classification-based approaches. Bowtie2 methods have previously been identified as highly effective for host read removal; as such we chose methods which facilitated comparison when sensitivity was adjusted and when combined with a classifier. We also wanted to investigate the performance of other methodologies which are widely used in the field, to assess their suitability against the popular alignment based methods. Three methods are based on the popular short-read aligner Bowtie2 (11), Bowtie2 standard is an implementation of the aligner with default parameters implemented within HOSTILE, which wraps Bowtie2 and performs the downstream removal of aligned reads (12). Bowtie2 VSL implements the “very-sensitive-local” parameter within Bowtie2 (11), increasing the sensitivity of Bowtie2 to more localised matches, mediated through a Kneaddata (13) wrapper with all other functionality switched off and read pairs being mapped in “unpaired” mode, hence treating each independently. We also implement Bowtie2 with a further Kraken2 (14) classification step for a 2-stage alignment followed by classification protocol. We also implemented the k-mer based method BMTagger as implemented within MetaWrap-QC (15), and the SRA-Human-Scrubber which is based on the SRA Taxonomy Analysis Tool (STAT) which NCBI recommends is run on read-level FASTQ data prior to submission and provides functionality to run the tool upon bioproject submission (16). For all methods, singleton reads were identified post-processing and removed, hence only paired reads which are not classified as human (either through alignment or classification) are retained for onward analysis.

**Table 1:** Details of the HRR methods investigated in this study.

| Method Name | Primary Removal Tool | Configurations | Wrapper | Human Reference(s) |
| --- | --- | --- | --- | --- |
| Bowtie2 Standard | Bowtie2 (v2.5.4) | Standard Bowtie2 parameters | HOSTILE (v1.1.0) | T2T-CHM13 and GRCh38 |
| Bowtie2 VSL | Bowtie2 (v2.5.4) | “Very-sensitive-local” parameter switched on and mapping performed on fastq pairs in “unpaired” mode | Kneaddata (v0.1.2) (all functionality except human read removal switched off) | T2T-CHM13 and GRCh38 |
| Bowtie2-Kraken2 | Bowtie2 (v2.5.4) for alignment, Kraken2 (v2.1.3) for subsequent classification | Standard parameters for Bowtie2 and Kraken2 | Nextflow v24.04.4 | Bowtie2: T2T-CHM13 and GRCh38<br>Kraken2: Standard-16GB database from Ben Langmead (https://benlangmead.github.io/aws-indexes/k2) |
| Metawrap-QC | BMTagger (v3.101) | Standard BMTagger configurations | Nextflow (https://github.com/sanger-pathogens/generate_mags) | T2T-CHM13 and GRCh38 |
| SRA Human Scrubber | SRA Human Scrubber (v2.2.1) | Standard SRA Human Scrubber Configurations | SRA Human Scrubber (v2.2.1) | 20240718 human database filter (https://ftp.ncbi.nlm.nih.gov/sra/dbs/human_filter/human_filter.db.20240718v2 ) |

We also sought to investigate the difference in performance when using different reference genomes, namely T2T-CHM13v2.0 (9) or GRCh38 (7). To this end, the three Bowtie2-based methods (Bowtie2 Standard, Bowtie2 VSL and Bowtie2-Kraken2) were run over samples sets twice, once with a T2T-CHM13 and again with a GRCh38 reference for comparison. For MetaWrap-QC, indexes were created using srprism (version 2.4.24-alpha) for each of the T2T-CHM13 and GRCh38 reference assemblies. For the SRA-scrubber, this approach was not applied, and we tested only the latest human filter database made available through the FTP (https://ftp.ncbi.nlm.nih.gov/sra/dbs/human_filter/), date stamped 2024/07/18.

Methods were orchestrated through a nextflow (17) pipeline (code found here: https://github.com/wtsi-npg/hrr_tools/tree/benchmarking_paper-2025), and runtime and memory usage statistics were obtained from nextflow trace files.

### Synthetic Mixture Titration Dataset Creation

We constructed a dataset to assess the performance of the different human read removal methods under investigation. This dataset contained real viral (RSV A and B, SARS-CoV-2 and Influenza A) and human reads mixed together in varying ratios, ensuring that reads from multiple human individuals with diverse genetic ancestry are present in the created read set mixtures. We selected 13 samples (7 from clinical sources, 6 synthetic control samples) sequenced through a bait capture procedure, and obtained all reads which aligned to one of three respiratory viruses: SARS-CoV-2, RSV A/B or Influenza A (see Supplementary Spreadsheet). Mapped paired-end reads were extracted, labelled for downstream identification and sorted with samtools v1.19. Human reads were obtained from a collection of 27 samples of different genetic ancestries from the 1000 genomes project (10), which was previously used in the evaluation of the HOSTILE HRR tool (12) (Supplementary Spreadsheet). This dataset was used as it was created to ensure a publicly available set of human sequences with diverse origins.

Viral and human reads were mixed in such a way as to ensure that the total number of human reads was sampled with identical frequency from each of the 27 human samples, ensuring that samples investigated comprised human reads from populations of different genetic ancestry in equal weight, thus allowing downstream investigation of differential ability to dehumanise samples taken from different human populations.

For each sample investigated, titration sets were created with viral:human proportions ranging from 1:9 - 9:1 via random sampling of viral and human read sets. Random sampling was performed in triplicate for each sample/human titre combination. This resulted in the generation of 351 paired-end fastqs representing the 13 viral samples, with 9 human read titres and 3 iterations of each (Fig. 1).

**Fig. 1:**
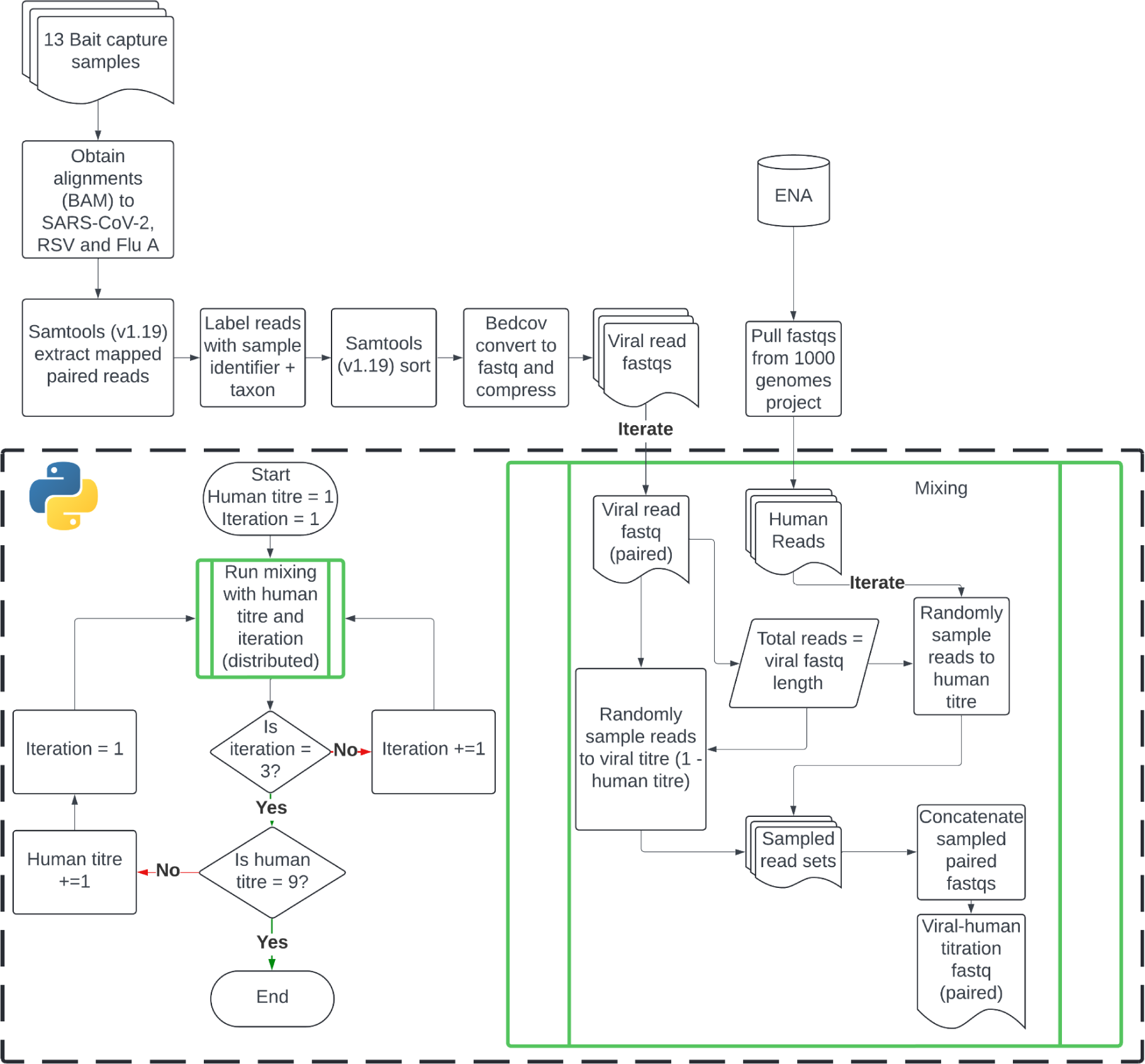
**Flowchart depicting the process for developing the synthetic mixture titration dataset.**

Read sets for the dataset are available through ENA, viral read sets for each sample can be obtained through the ERP169280 project, titration mixtures can be obtained through the ERP169281 project and titration mixtures processed through the various HRR methods can be obtained through ERP169282. A full mapping of all samples can be found in supplemental materials.

### Calculation of Statistics

After titration sets were run through each human read removal method, read names were extracted from both the removed and retained read sets to obtain the status of all reads in the investigation. Binary classifiers were calculated for each titration sample; we defined a “positive” case as a read removed by a human read removal method and established the truth status depending on whether the read was incorporated into the dataset from viral or human origin (e.g. a read removed from the dataset by a HRR method which originated from a human sample will be labelled as a true positive).

These binary classifiers were subsequently used to classify performance metrics, this was split into “all-round” metrics such as accuracy, Matthew’s Correlation Coefficient (MCC) and F1 scores, and more specific scores such as Precision, Sensitivity (also known as True Positive Rate or Recall) and Specificity (also known as True Negative Rate).

### Investigation of False Negative Read Sets

False negative reads were put through taxonomic classification by Kraken2 with default settings (14), using the maxikraken database supplied by the Loman group for the Mock Communities project (url: https://lomanlab.github.io/mockcommunity/mc_databases.html) (18). Percentages were extracted from Kraken2 reports and in-house code was used to parse reports and to merge counts across samples retaining hierarchies.

### Evaluation of Consensus Fasta Generation

The human-read-removed titration datasets 1, 5, and 9 were trimmed and filtered using Trimmomatic (version 0.39), read sets were then taxonomically classified using Kraken2 (version 2.1.3) with a custom database optimised for reference selection consisting of the contents of Viral RefSeq (19), with additional influenza and RSV genomes obtained from NCBI. To improve the species selection, a bespoke taxonomy was applied within the database which included 1) pruning sub-species leaf nodes for non-influenza/RSV genomes, 2) reorganising RSV genomes into a hierarchical sub-structure for types A and B, and 3) reorganising influenza genomes to effectively bin reads by influenza genome segments. For segments 4 and 6, an additional hierarchical layer was introduced to distinguish between HA and NA types. Viruses with more than 100 assigned reads were selected as references for mapping with BWA (version 0.7.17) and consensus sequences were generated with iVar (version 1.4.3), calling bases with coverage equal to or greater than 10X (pipeline code here: https://github.com/genomic-surveillance/rvi-viral-lens, manuscript under construction).

The percentage of sequence coverage against various selected reference sequences was assessed as the proportion of positions in the alignment covered to at least 10X, and was subsequently used to perform a Principal Component Analysis (PCA) followed by a t-distributed Stochastic Neighbor Embedding (t-SNE, perplexity 30, number of iteration 5000) for visualisation. A PERMANOVA (Permutational Multivariate Analysis of Variance) was performed to assess the statistical significance of differences between groups within the dataset.

### Evaluation of External Metagenomics Dataset

We selected a publicly available shotgun metagenomics study of the nasal microbiota to benchmark our method, accession number CRA006819 deposited on Genome Sequence Archive (GSA). To date, this is the largest publicly available shotgun metagenomic dataset for the nasal microbiota. Before the data was uploaded, the authors removed human reads with Bowtie2 using the human genome GRCh38 as the reference (20). Raw FASTQ files were downloaded from the GSA. The human read removal pipelines, as described in Table 1, were applied to the dataset. The microbiota taxonomic composition was determined using sylph 0.6.1 (21) with GTDB release 220 (22) sketched at c200.

Statistical analysis was performed in R version 4.4.2 (23). Alpha diversity was calculated with the vegan package version 2.6-10 (24) with the functions specnumber and diversity. Pielou’s evenness was calculated using the function diversity found within the microbiome package version 1.28.0 (25). Kruskal-Wallis test and pairwise Wilcox test were used to test for statistical significance. For beta diversity, Bray-Curtis dissimilarity and Jaccard Index were calculated using vegdist function found within vegan. A pairwise t-test was used for statistical testing. P-values were adjusted with the Benjamini-Hochberg procedure. Pairwise PERMANOVA was calculated with function pairwise.adonis2 from the package pairwiseAdonis (26). Visualisation of the average nucleotide identity was done with ggalluvial (27). MaAsLin 2 v1.20.0 was used to test for microbial association (28). The fixed effect measured is the method used for human read removal while the random effect was the sample. The raw dataset (Bowtie2 Standard with GRCh38) was set as the reference. The default parameters were used. To visualise the difference between the methods, a PCoA was calculated using the pcoa package from ape v5.8-1 (29) and the dbrda functions found within the vegan package. The function varpart was used to calculate the partition of variation in the data.

To assess the ability to determine human sample sex from uploaded metagenomic data, a subset of 10 samples (M=5, F=5) were used. We compared pre-human read removal samples (uploaded to GSA) with post-human read removal samples (GSA samples processed through SRA human scrubber, Bowtie2 VSL with a GRCh38 reference genome, and Bowtie2 VSL with a T2T-CHM13v2.0 reference genome). Post-human read removal samples were re-aligned to the T2T-CHM13v2.0 reference using Bowtie2 version 2.5.1 and the ‘--very-sensitive-local’ option to identify retained human reads. Information about the proportion of reads mapping to the reference genome was gathered using samtools1.17.

Mean read depth values were visualised using the ‘pseudo_log_trans()’ function from the ‘scales’ R package version 1.3.0.

## Results

### Comparison of Methodology Performance

We sought to identify the best practice for human read removal from microbial metagenomics short read data (shotgun and bait capture). To do this, we investigated the performance of multiple Human Read Removal (HRR) approaches (see details of selection in Methods and Table 1).

All methodologies tested showed good performance in human read removal across the synthetic titration benchmark, with the minimum Matthew’s Correlation Coefficient (MCC) score at any human titre for any method of 0.94 (Fig. 2A). As the human titre is increased, accuracy and MCC show a pattern of decay across all methods at similar rates, with no method converging or crossing across human titres. This decay is not shown for the F1 score (the harmonic mean of precision and recall), indicating that this performance decay was driven mainly by the increasing number of false negatives (Fig. S1), which is not a component of the F1 score.

**Fig. 2:**
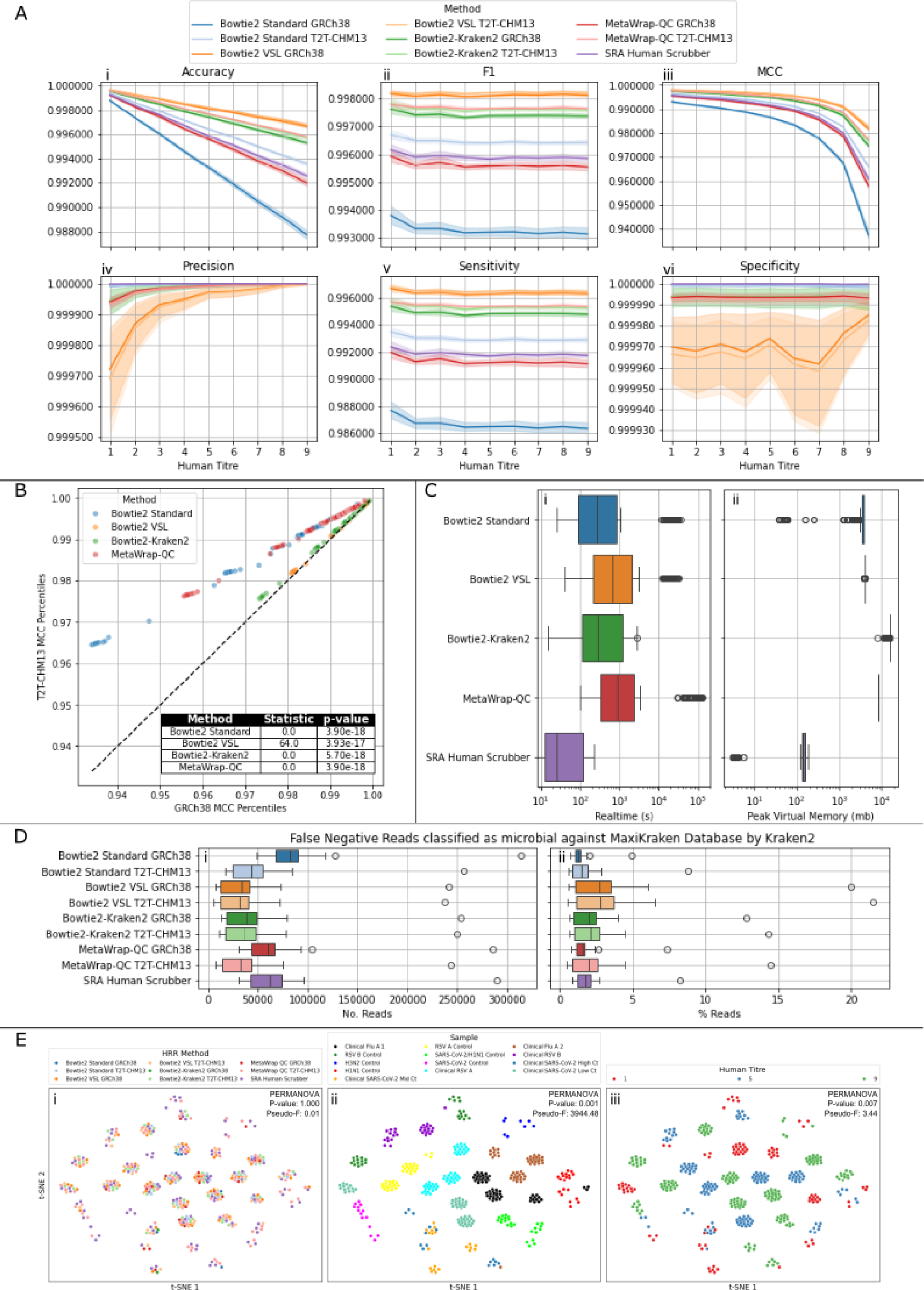
Comparison of Human Read Removal Methods across the Synthetic Titration Dataset. A) Performance metrics across the dataset, top row (i-iii) shows overall classification performance: Accuracy (i) reflects the proportion of correctly classified reads but can be misleading with class imbalance; F1 Score (ii) balances Precision and Recall, making it useful when false positives and false negatives have different consequences; Matthews Correlation Coefficient (MCC, iii) provides a more robust measure by incorporating all elements of the confusion matrix. The bottom row presents class-specific metrics: Precision (iv) indicates the proportion of correctly identified human reads among those classified as human, Sensitivity (v) shows how effectively human reads are removed, and Specificity (vi) measures how well viral reads are retained. Lines indicate the mean across all samples and iterations at each human titre, with shading representing the 95% confidence interval. Different y-axis scales between plots should be considered when interpreting values. B) Percentile-percentile plot showing the differences in Matthew’s Correlation Coefficient (MCC) distributions between methods run with the GRCh38 and T2T-CHM13 human assemblies as reference sequences. Table shows the output of Wilcoxon signed rank test for each method between the GRCh38 and T2T-CHM13 distributions. C) Boxplot showing differences in run time and resource usage between the different methodologies. (i) realtime for process completion in seconds. (ii) Peak Virtual Memory used in the completion of the process in megabytes. D) Boxplots showing distributions of false negative (reads derived from human sources retained by HRR methods) reads which are classified as microbial (i.e. neither human nor unclassified) by kraken2 against the Maxikraken database across the synthetic titration dataset. (i) no. false negative reads classified as microbial, (ii) percentage thereof. E) t-SNE embeddings of consensus sequence coverage after HRR and comparing different human read titres. Points coloured by HRR method (i), Sample (ii) and human titre (iii), with PERMANOVA P-value and Pseudo-F values for each grouping indicated in each plot.

Across the methodologies investigated, the best all-around performing methods were those utilising Bowtie2 with the “very-sensitive-local” (VSL) configuration, as indicated by the clear difference in all-around performance showcased through Accuracy, F1 and MCC scores. The Bowtie2-Kraken2 methods had the next best performance, with the poorest performance exhibited by the Bowtie2 Standard methodology with the GRCh38 human reference (Fig. 2A).

Investigation of specific components of the overall performance indicated that the increase in sensitivity conferred by the Bowtie2 VSL methodology also resulted in a very small drop in specificity (true negative rate, the proportion of actual negatives correctly identified by a test), indicating that this method was less specific than other methods, and more prone to calling false positives than other methods. This difference is marginal, with a reduction in specificity of ∼0.00003 from the highest specificity method (SRA Human Scruber), and with a median false positive count of 5 reads with a maximum of 375 (the Bowtie-Kraken2 method has a higher maximum False Positive rate, see Fig. S1 and Table S1). This marginal reduction in specificity does not overcome the larger increase in sensitivity (true positive rate, the proportion of actual positives identified by a test), contributing to its greater performance in the overall metrics compared to other methods.

Overall, the best-performing method, Bowtie2 VSL T2T-CHM13, exhibited an approximately 3.8 fold reduction in the number of false negative reads (i.e. human reads incorrectly retained by the method) compared to the worst-performing method, Bowtie2 Standard GRCh38. It also exhibited an approximately 1.3 fold decrease compared to the next-best T2T-CHM13 method in Bowtie2-Kraken2.

The differences in human reference sequence appeared to have a different effect according to the different HRR methodologies (Fig. 2B). All methodologies showed a significant improvement in HRR when the T2T-CHM13 reference sequence is applied according to the MCC scores (Wilcoxon signed rank test p-value < 0.05). The extent of this was variable between methods, with Bowtie2 Standard and MetaWrap-QC showing the largest difference between the two methods; with a much smaller difference for the Bowtie2 VSL and Bowtie2-Kraken2 methods (Bowtie2 VSL also shows a higher test statistic, which suggests that the rank scores vary more within the dataset and as such there is less distinction between the two populations being compared. Fig. 2B). This indicates that the sensitivity of the method employed is a more significant component of HRR performance than the reference sequence used and that the more sensitive methods (Bowtie2 VSL and Bowtie2-Kraken2) are less likely to be improved further by improvement in human genome assemblies reducing the maintenance burden of these methods.

We also examined resource usage across the tested methods (Fig. 2C). The VSL parameterisation had a significant impact on overall runtime (Wilcoxon rank-sum test, *p* < 0.05), resulting in a median increase of 6.42 minutes compared with the standard Bowtie2 implementation. However, memory consumption was similar between Bowtie2 VSL and the standard Bowtie2, while Bowtie2-Kraken2 and MetaWrap exhibited higher memory usage. In contrast, the SRA Human Scrubber was markedly more efficient than the other methods, with substantially lower runtime and peak memory consumption.

A key focus for HRR methodologies is the effective removal of host reads, therefore we focused the analysis on the composition of reads which were recorded as false negatives (FNs, reads arising from human samples which were not removed by a given methodology). As these FNs all arose from samples from the 1000 genomes project, we investigated the extent to which these retained reads were true human reads rather than probable microbial and/or other contaminants in the 1000 genomes samples. To this effect, FN read sets were run through Kraken2 against the “maxikraken” database (see Methods) and their compositions were compared between methods.

As a percentage of FN reads, the Bowtie2 VSL methods exhibit significantly (Wilcoxon rank p-value < 0.05) increased microbial reads compared with other methods, indicating that human and unclassified reads make up for a much smaller component of these read sets resulting from these methods (Fig. 2D(i)). At the same time, we can see that the overall number of microbial reads retained by the VSL methods is decreased for these methods compared with other methods (Fig. 2D(ii)). Taken together, this provides a stronger indication that these methods have a greater propensity to deplete potential human reads from the dataset, however, with greater potential to additionally deplete some real microbial reads. FN reads which are either classified as human or that remain unclassified after Kraken2 processing did not exhibit bias towards certain genetic ancestries (Fig. S2). The HRR method employed appears to be a much stronger component of read retention than that of human genetic ancestry.

To assess the impact of different HRR methods on viral consensus genome FASTA generation, we investigated genome coverage patterns across different viral species with human titres 1, 5, and 9. Dimensionality reduction revealed that clustering patterns of consensus fasta coverage were primarily driven by the sample and differences in human titres (Fig. 2E(i) and (ii)), with little to no contribution from the HRR methods (Fig. 2E(iii)). In line with this, PERMANOVA showed no significant grouping based on HRR methods (Fig. 2E(iii)). Collectively, these findings suggest that the HRR process had minimal influence on the overall viral consensus genome qualiity.

### Evaluation of external metagenomic data

To evaluate the effect of HRR methods, we applied the approach to a publicly available shotgun metagenomic study. Human read removal had already been performed by the authors using Bowtie2 Standard GRCh38, prior to the dataset being uploaded into the public domain. We observed a significant difference in richness (number of observed bacterial species), Shannon diversity (diversity of the community taking into account the number of observed bacterial species and relative abundance), Inverse Simpson (similar to Shannon diversity but emphasises on community evenness), and Pielou’s evenness (how evenly distributed the species are in a community) between raw to SRA Human Scrubber, Bowtie2 VSL GRCh38, and Bowtie2 VSL T2T-CHM13 (Fig. 3A). However, no significant differences were found between Bowtie2 VSL GRCh38 and T2T-CHM13. Bowtie2 VSL, regardless of reference, reduced detectable bacterial species from 31 in raw to 11 (Table S2). Shannon diversity and Inverse Simpson also decreased. Notably, Bowtie2 VSL increased evenness, suggesting preferential removal of less abundant species.

**Fig. 3:**
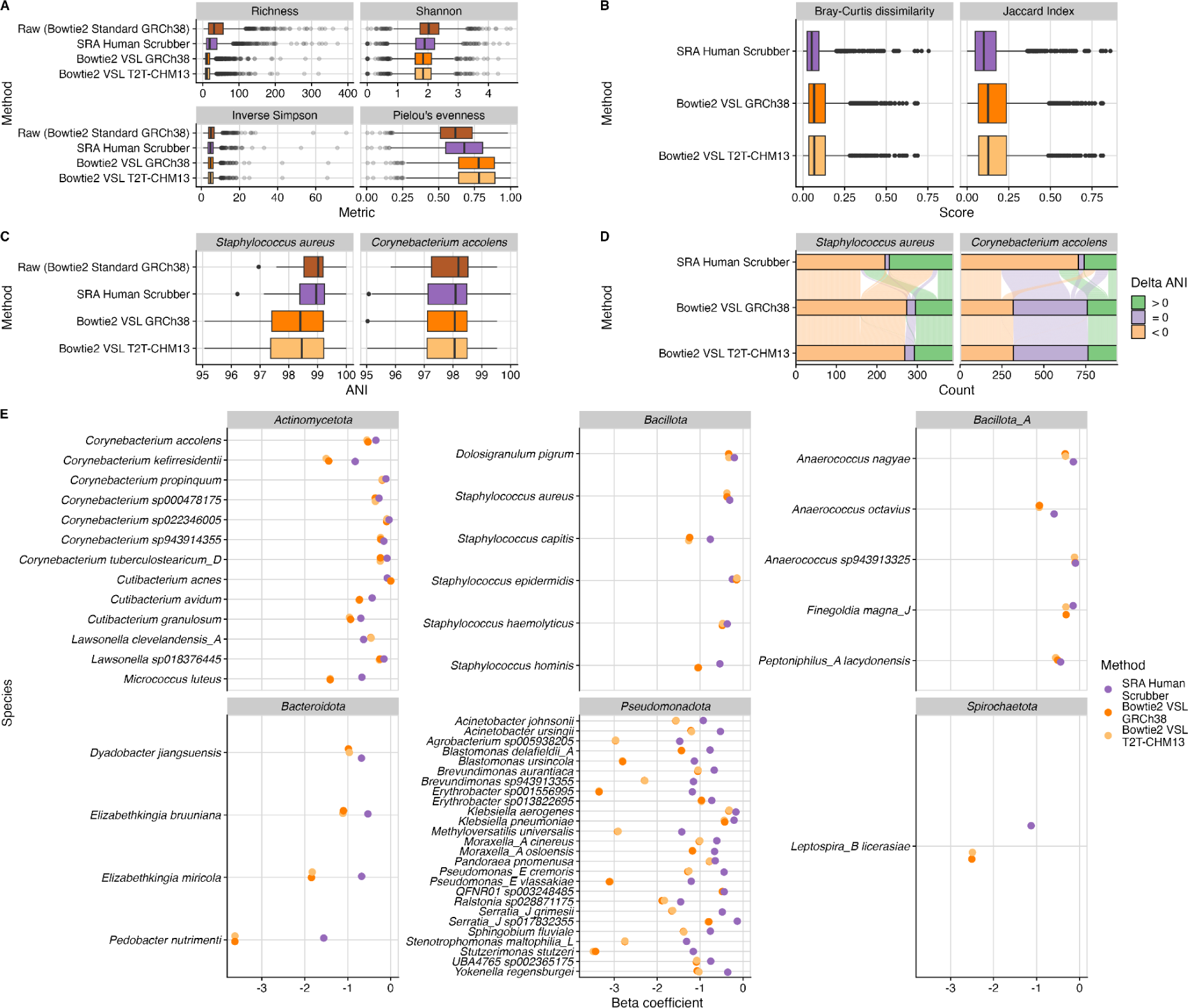
Comparison of metagenomic outputs arising from metagenomics analysis post implementation of HRR methods. (A) Alpha diversity comparison between three HRR methods and a standard HRR approach, Bowtie2 standard GRCh38. (B) Bray-Curtis dissimilarity and Jaccard index score of three HRR methods against the raw. (C) Visualisation of the average nucleotide identity (ANI) of two species commonly found in the nasal microbiota. (D) Delta ANI score of three HRR methods against the raw. (E) A differential analysis of HRR methods compared to the raw showing the depletion of bacterial species.

We evaluated the effect of HRR methods on beta diversity, which measures differences in microbial community composition between samples, using Bray-Curtis dissimilarity on relative abundance. Significant differences were observed between raw and SRA Human Scrubber (R² = 0.0009, p = 0.003), Bowtie2 VSL GRCh38 (R² = 0.00357, p = 0.001), and Bowtie2 VSL T2T-CHM13 (R² = 0.00364, p = 0.001). However, ordination did not show clear separation between clusters (Fig. S3A). Variation partition analysis attributed 0.36% of variation to HRR methods and 97.26% to samples (Fig. S4). After accounting for sample variation, a slight separation between methods was observed (Fig. S3B), with Bowtie2 VSL clustering together regardless of the reference genome used. Bray-Curtis dissimilarity and Jaccard index scores were used to quantify differences (Fig. 3B). SRA Human Scrubber had median scores of 0.05 (Bray-Curtis) and 0.1 (Jaccard). Bowtie2 VSL showed similar median scores across references: Bray-Curtis 0.067 (GRCh38) and 0.068 (T2T-CHM13); Jaccard 0.125 (GRCh38) and 0.1268 (T2T-CHM13).

Next, we compared the determined average nucleotide identity (ANI) of two common species found in the nasal microbiota; *Staphylococcus aureus* and *Corynebacterium accolens* across the HRR methods (Fig. 3C). We hypothesised that ANI increases when human reads misclassified as bacterial are removed, while ANI decreases when bacterial reads are misclassified and removed (Fig. S5). *S. aureus* ANI scores differed significantly between the raw dataset and Bowtie2 VSL (GRCh38 and T2T-CHM13) but not between raw and SRA Human Scrubber. No significant difference was observed between Bowtie2 VSL GRCh38 and T2T-CHM13, suggesting Bowtie2 VSL itself, rather than the reference genome, drives the difference. SRA Human Scrubber improved *S. aureus* ANI scores, whereas Bowtie2 VSL reduced them (Fig. 3D) suggesting *S. aureus* reads are removed by Bowtie2 VSL. For *C. accolens*, ANI scores differed significantly between raw and both HRR methods but not between SRA Human Scrubber and Bowtie2 VSL, indicating similar performance.

Notably, Bowtie2 VSL performed better than SRA Human Scrubber at maintaining or improving *C. accolens* ANI scores, suggesting that the extent of misclassified bacterial reads may vary by species.

To identify species removed by different HRR methods, we performed differential abundance analysis using MaAsLin2. Of the 3,115 species detected, 55 were depleted due to HRR (Fig. 3E), primarily from the phyla *Pseudomonadota* and *Actinomycetota*. Bowtie2 VSL GRCh38 and T2T-CHM13 had similar beta coefficients (mean delta: 0.0001). SRA Human Scrubber had a lower beta coefficient than Bowtie2 VSL, with a mean delta of 0.58 for both GRCh38 and T2T-CHM13 (Table S3).

In addition to quantifying the impact of human read removal methodologies on microbiome diversity, we assessed the post-HRR samples for the presence of residual human reads. The aim of this analysis was to determine the level of human genetic information retained in post-HRR samples that could potentially be used to re-identify individuals. Reads were retained across autosomes and sex chromosomes in samples processed with the Bowtie2 standard GRCh38 reference method and the SRA human scrubber method; Bowtie2 VSL with a GRCh38 reference genome retained notably fewer reads than both these methods (Fig. 4A). Bowtie2 VSL with T2T-CHM13v2.0 reference genome did not detect retained reads, which was expected as the same reference genomes were used for initial HRR and subsequent re-alignment.

**Fig. 4:**
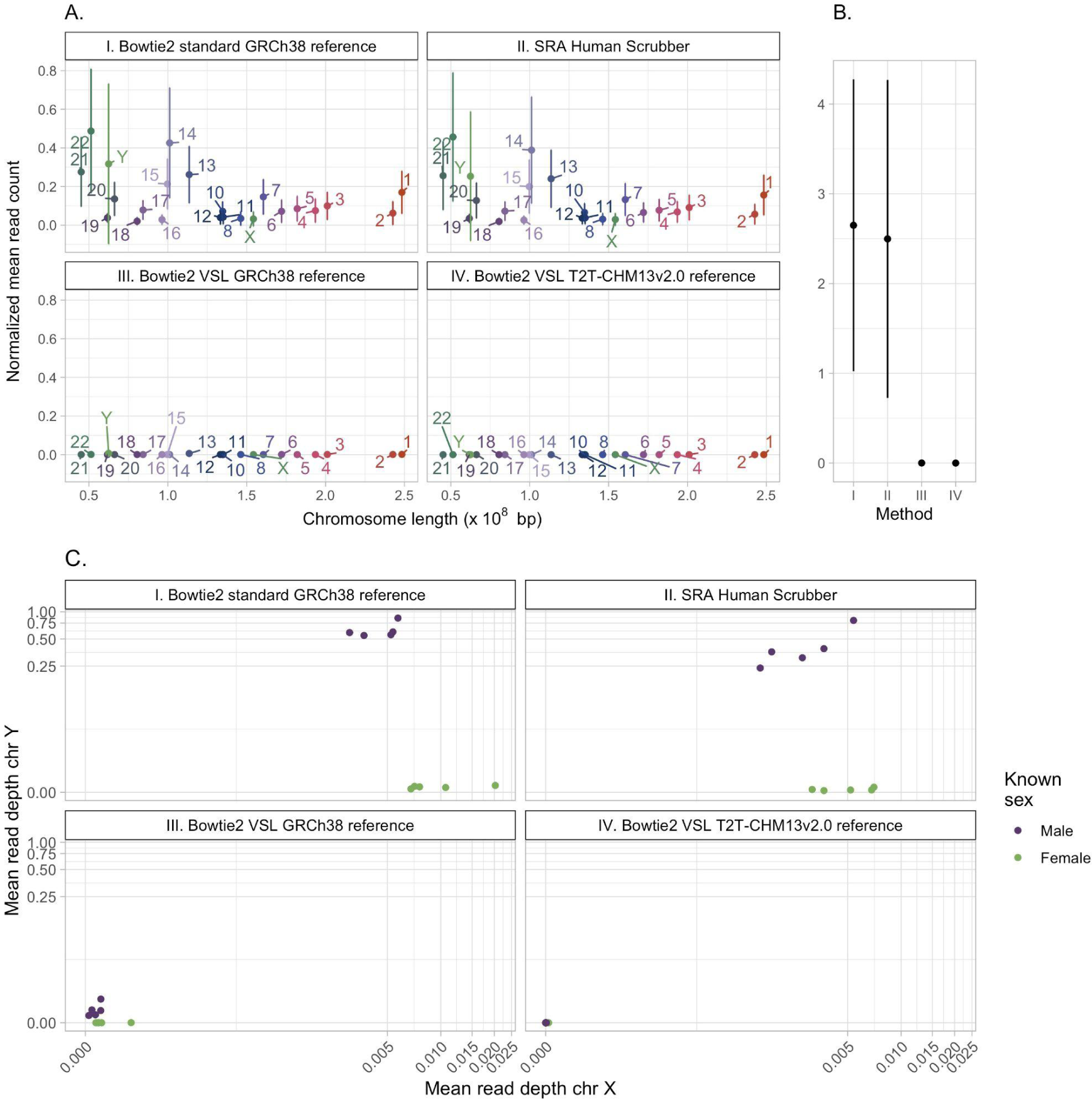
Post human read removal analysis of public human nasal microbiome data. (A) Normalized mean read count when using (I) Bowtie2 standard with a GRCh38 reference (data as released by Ju et al., (20)), (II) SRA human scrubber, (III) Bowtie2 VSL with a GRCh38 reference, and (IV) Bowtie2 VSL with a T2T-CHM13v2.0 reference. Normalized mean read count on chromosome 9. (**C**) Plots compare the pseudo-log mean read depth of chromosome X and Y post human read removal using (I) Bowtie2 standard with a GRCh38 reference, (II) SRA human scrubber, (III) Bowtie2 VSL with a GRCh38 reference, and (IV) Bowtie2 VSL with a T2T-CHM13v2.0 reference.

Improvements made by the T2T-CHM13v2.0 reference genome include the resolution of highly repetitive regions, such as the human satellite array on chromosome 9, a region which has previously been of low mappability (30). It is notable that a larger number of human reads were retained on chromosome 9 by the Bowtie2 standard or SRA human scrubber methods (Fig. 4B). This demonstrates the importance of using a high-quality reference genome for human read removal.

We earlier highlighted the ethical concerns that surround the ability to re-identify an individual from human reads retained in microbiome data. This was previously tested in a cohort of individuals with gut microbiome samples and corresponding genomic samples (4). Without access to human biobank data associated with the nasal microbiome samples we are unable to fully test the ability to correctly match microbiome samples to human genetic samples.

As an approximation of re-identification, we performed a sex determination analysis of samples post-HRR (Fig. 4C). Sex determination was possible for all samples in the publicly uploaded nasal microbiome shotgun data (mean Y chromosome read depth in male samples, rd = 0.692), and remained possible after HRR with SRA human scrubber (rd = 0.420) or Bowtie2 VSL with a GRCh38 reference genome (rd = 0.00673), although with diminishing confidence. SRA scrubber and Bowtie2 VSL GRCh38 methodologies appear to be proficient in removing X chromosome reads, but retain a larger proportion of Y chromosome reads; Bowtie2 VSL GRCh38 to a much lesser extent, reflecting the improved performance of the Bowtie2 VSL method. The impact of reference genome is highlighted by the differences observed between Bowtie2 VSL using the GRCh38 reference genome and the T2T-CHM13v2.0 reference genome; reflecting the improvements made by the T2T-CHM13v2.0 reference genome, including the more accurate representation of the Y chromosome included in the T2T-CHM13v2.0 reference sequence (31).

## Discussion

This investigation demonstrates that the “very-sensitive-local” (VSL) mode in Bowtie2 significantly enhances human read removal (HRR) performance by increasing alignment sensitivity at the cost of longer runtime (Table 2). Unlike the default mode, VSL enables local alignment, allowing soft-clipping of non-matching bases at read ends, making it more tolerant to sequencing errors. The seed length is reduced from 22 to 20 bases, increasing sensitivity but also computational time, as shorter seeds match more frequently across the genome. Additionally, VSL increases re-seeding attempts from 2 to 3, improving alignment chances when the initial seed fails. The seed interval factor is lowered from 1.0 to 0.5, resulting in denser seed placement, which enhances alignment probability but requires more processing. Finally, the allowed backtracking is increased, allowing Bowtie2 to explore more alternative alignment paths. Collectively, these modifications increase the likelihood of mapping reads to the human reference but require additional computational resources.

**Table 2:** Differences in Bowtie2 parameters between the Default and the Very Sensitive Local modes.

| Mode | Alignment Type | Seed Length | Backtracking | Re-Seeding | Seed interval factor |
| --- | --- | --- | --- | --- | --- |
| Default | End-to-end | 22 | 15 | 2 | 1 |
| Very Sensitive Local | Local | 20 | 20 | 3 | 0.5 |

Another key difference in the VSL methodology we have implemented is that reads are aligned to the reference by Bowtie2 in “unpaired” mode. This has the ability to confer several further sensitivity gains aside from the implementation of the “very-sensitive-local” mode alone, including the removal of constraints around insert size restrictions, proper pairing of reads, greater sensitivity to partial matches (e.g. where one mate appears human and the other microbial) and potential handling of structural variants within human genomes which may disrupt insert sizes. Furthermore, the removal of the requirement to compute pair restraints in unpaired mode likely reduces resource requirements hence partly compensating for the increased resource usage of the “very-sensitive-local” modality.

The combination of the parameter changes, allowance of soft-clipping and removal of pairing constraints has a clear positive effect on the sensitivity of the Bowtie2 VSL methodology; however, this increase in sensitivity has obvious potential drawbacks, including additional computation time and a decrease in specificity. Indeed we did observe a marginal decrease in specificity for the Bowtie2 VSL method over other methodologies. However, when comparing the difference between the top T2T-CHM13 methods (Bowtie2 VSL and MetaWrapQC), the difference in median sensitivity is approximately 191 times that of specificity. This is reflected in the fact that the overall metrics (accuracy, F1 and MCC) are in agreement that Bowtie2 VSL has the most optimal overall performance. Additionally, this decrease in specificity is reflected only in the removal of 5 viral reads in the median case, with 375 (out of a total of 8,005,608 reads) in the worst case (the worst case across the whole dataset is 429 reads from the Bowtie2-Kraken2 method out of a total of 8,563,378).

Bowtie2 VSL has an effect on microbial composition, but the effect size is small. After HRR by Bowtie2 VSL, rarer species are excluded, leading to a more even community. HRR can introduce batch effects, and if multiple HRR methods are applied unevenly, they may bias the dataset. Authors should clearly report each step to ensure reproducibility. Bowtie2 VSL can also misclassify bacterial reads with GC content similar to humans (40.9%). This effect was more pronounced in *S. aureus* (GC 32.8%) than *C. accolens* (GC 59%), possibly due to Sylph’s k-mer-based microbiota quantification, which is more sensitive to small changes in read composition (ref for Sylph).

Additionally, it is clear that implementation of Bowtie2 VSL in combination with the T2T-CHM13v2.0 reference genome is the most effective for removing host-contaminating reads; consequently providing the greatest security against identification, demonstrated by the reduced ability to identify the sex of the individual host. As such, although a small effect on microbial diversity is observed, this is outweighed by the reduction in the ability to identify human individuals.

More broadly, our investigation indicates that the selection of a human read removal method and its configurations likely depends mostly on scalability and robustness. We observe that overall metrics such as Accuracy and MCC show a reduction in overall performance as the proportion of human reads in the titration set is increased. This is likely driven by two main factors: the presence of more human reads translating into an increased chance of misidentification, and the inclusion of more true microbial contaminants from the 1000 genomes samples. Moreover, as the sensitivity remained relatively constant through the titration series for all methods, there is no indication that methods should be chosen on the basis of likely human titre. We also found no evidence of bias based on human genetic ancestry in the removal of human reads, with methodology seemingly the most important component. Therefore, without questions of scalability, the implementation of Bowtie2 VSL T2T-CHM13v2.0 would appear to be optimal.

When we consider scalability, however, we must take into consideration that Bowtie2 VSL shows a median time to completion of around 10.91 minutes, which is an increase of around 6.42 minutes per sample longer than the median time for the Bowtie2 Standard configuration. While this might be appropriate for users without time constraints or those with large computational resources allowing them to highly parallelise the process, these timescales can become burdensome. The memory profile, however, was fairly similar between the Bowtie2 standard and VSL methods, with the addition of the Kraken2 classification step increasing the memory usage by around 4-fold.

By far the most efficient tool by both runtime and memory usage was the SRA Human Scrubber, which uses the fast STAT methodology combined with a pre-indexed version of the human genome covering most regions of interest (16). This allows the Scrubber to take a more lightweight approach to human read removal, performing a fast scan over a large proportion of the identifiable human genome rather than a full scale alignment. Our investigation indicates that this approach results in the highest specificity of any method in the benchmark, with a maximum of only 2 false positive calls across the entire benchmark.

However, we also observe that the scrubber’s sensitivity is substantially lower than that of the best-performing methodology, Bowtie2 VSL, and is also lower than that of the Bowtie2 Standard implementation with the T2T-CHM13v2.0 reference genome. This indicates that a greater proportion of human reads will make up these samples, around 2,592 more reads than Bowtie2 VSL in the median case, which could have downstream effects on identifiability of human sequences, the composition of consensus sequences and on downstream runtime. Our investigation indicates that coverage of consensus sequences does not appear to be affected by the use of either the SRA Scrubber or the Bowtie2 VSL methodology compared with other methods.

Overall, our investigation indicates that all the methods tested here are likely to be suitable for human read removal in a variety of experiments, all the methods show high sensitivity and specificity. For the greatest reduction in potential risk for submission to public archives, the Bowtie2 VSL is most likely to remove the largest proportion of human reads and therefore reduce downstream risk to the greatest extent. There are two potential caveats to this: the potential removal of a larger number of microbial reads and the increased resource usage of the higher sensitivity bowtie2 methodology. In terms of incorrect removal, this analysis indicates that this effect, while present, is minimal, and outweighed by the positive effect of increased removal of host reads and subsequent reduction in identification risk. For resource usage, tools such as the SRA-human-scrubber are available, however, this investigation indicates that usage of these tools incurs a compromise on downstream data security. Instead, further developments could be made to implement methods which match the sensitivity of Bowtie2 VSL while reducing resource usage, a potential example may be implementation of HISAT2 which may be able to offer similar sensitivity but reduce resource consumption through it’s usage of a hierarchical index, improved memory consumption and cache optimisation (32).

Finally, when using a reference or reference k-mer database for alignment-based removal, the use of the latest T2T-CHM13v2.0 reference is recommended, however using more sensitive methods such as Bowtie2 VSL can help to ensure that your method remains robust to improvements in human genome assemblies without creating new vulnerabilities.

## Data Availability

Read sets for the dataset are available through ENA, viral read sets for each sample can be obtained through the ERP169280 project, titration mixtures can be obtained through the ERP169281 project and titration mixtures processed through the various HRR methods can be obtained through ERP169282. A full mapping of all samples is provided in a separate spreadsheet alongside this paper.

## Acknowledgements

The authors would like to thank Antonio Marinho, Frank Schwach, Katherine Figueroa, Thomas Maddison, Simon Suddaby, Marissa Knoll and Diego Teixeira for contributions to discussions of the work. Additionally, we would like to thank Katie Bellis, Steve Walton, Quan Lin and Jayvant Desale for their support in submission of data to ENA, and to Jaime Tovar Corona, Salih Tuna, Waleed Osman and Kevin Lewis for precursor evaluations leading to the work. This work has been supported by the Wellcome Trust [220540/Z/20/A].

## Author Contributions

● Conceptualisation: Matthew Forbes, David Jackson, Kevin Howe, Ewan Harrison
● Methodology: Matthew Forbes, Ewan Harrison, Kevin Howe, Duncan Y. K. Ng, Róisín M. Boggan, Bruhad Dave
● Software: Jillian Durham, Oliver Lorenz
● Validation: Matthew Forbes, Duncan Y. K. Ng, Róisín M. Boggan, Andrea Frick-Kretschmer, Florent Lassalle, Carol Scott, Josef Wagner, David Jackson
● Formal analysis: Matthew Forbes, Duncan Y. K. Ng, Róisín M. Boggan, Andrea Frick-Kretschmer
● Investigation: Matthew Forbes, Duncan Y. K. Ng, Róisín M. Boggan, Andrea Frick-Kretschmer, Jillian Durham
● Data Curation: Matthew Forbes, Duncan Y. K. Ng
● Writing - Original Draft: Matthew Forbes, Duncan Y. K. Ng, Róisín M. Boggan, Andrea Frick-Kretschmer
● Writing - Review and Editing: Matthew Forbes, Duncan Y. K. Ng, Róisín M. Boggan, Andrea Frick-Kretschmer, Jillian Durham, Ewan Harrison
● Visualisation: Matthew Forbes, Duncan Y. K. Ng, Róisín M. Boggan, Andrea Frick-Kretschmer
● Supervision: Ewan Harrison, Kevin Howe, David Jackson
● Project administration: Adrianne Lignes, Fernanda Noaves
● Funding acquisition: Ewan Harrison

## Supplementary

**Fig. S1:**
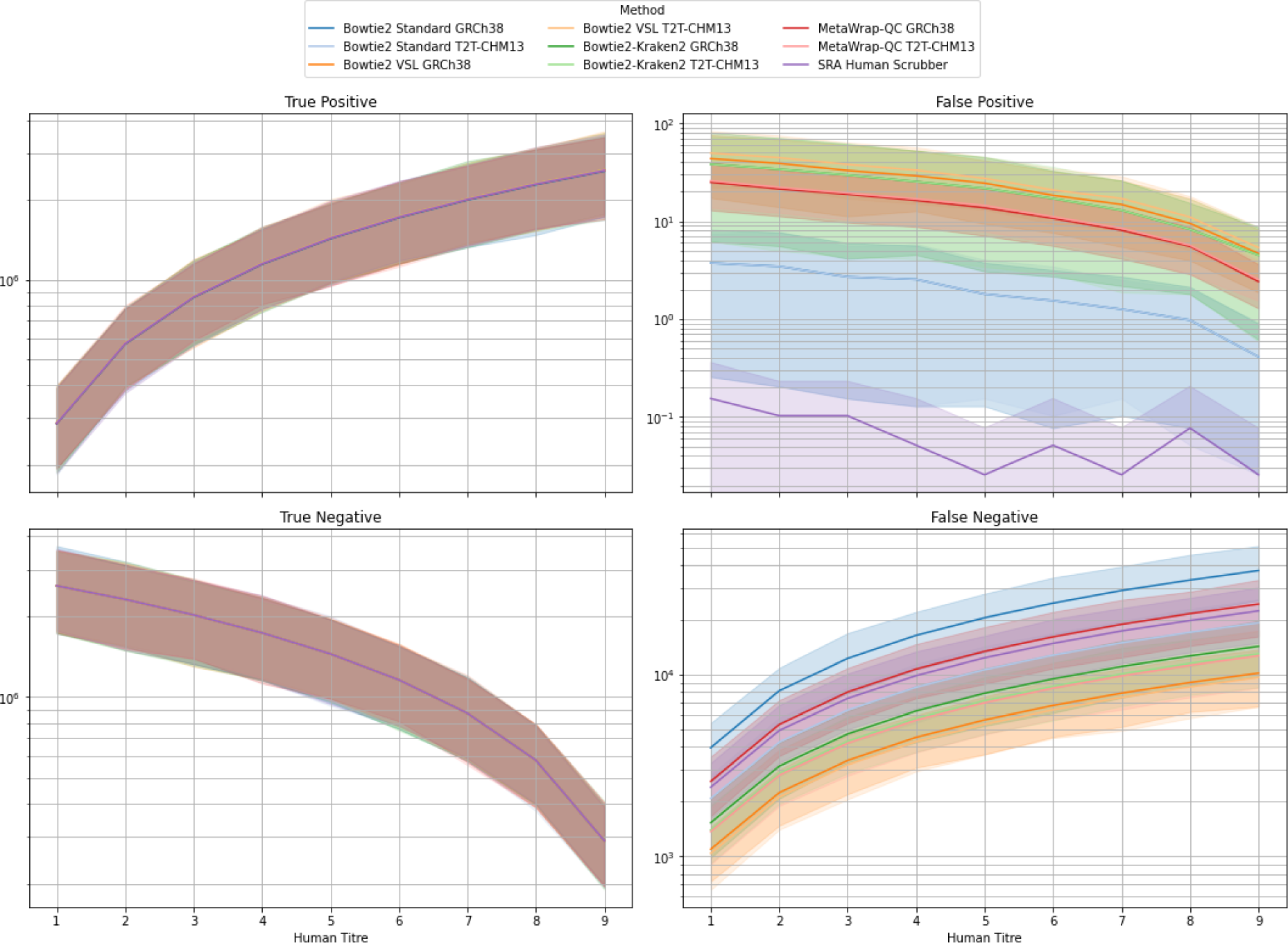
Comparison of binary classifiers with regard to the Human Read Removal methods over the synthetic titration dataset. Lines show the mean value across all samples/iterations at a given human titre, shading shows the 95% confidence interval around the mean. Note the different scales on the y-axis between plots when interpreting.

**Fig. S2:**
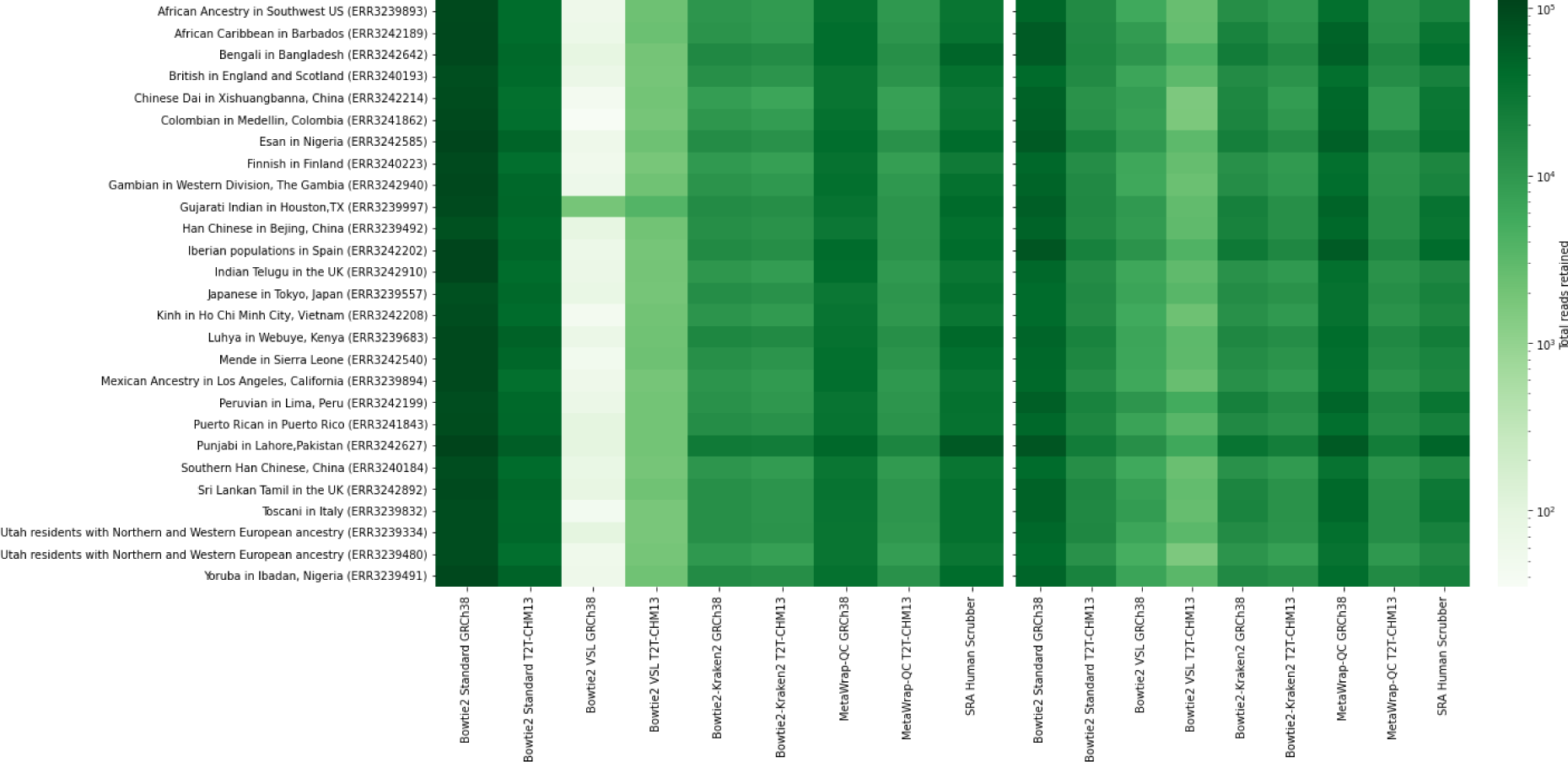
Heatmaps showing the number of false negative reads (reads derived from human sources retained by HRR methods) which were classified by Kraken 2 as human (left plot) or were unclassified (right plot) per genetic ancestry.

**Fig. S3:**
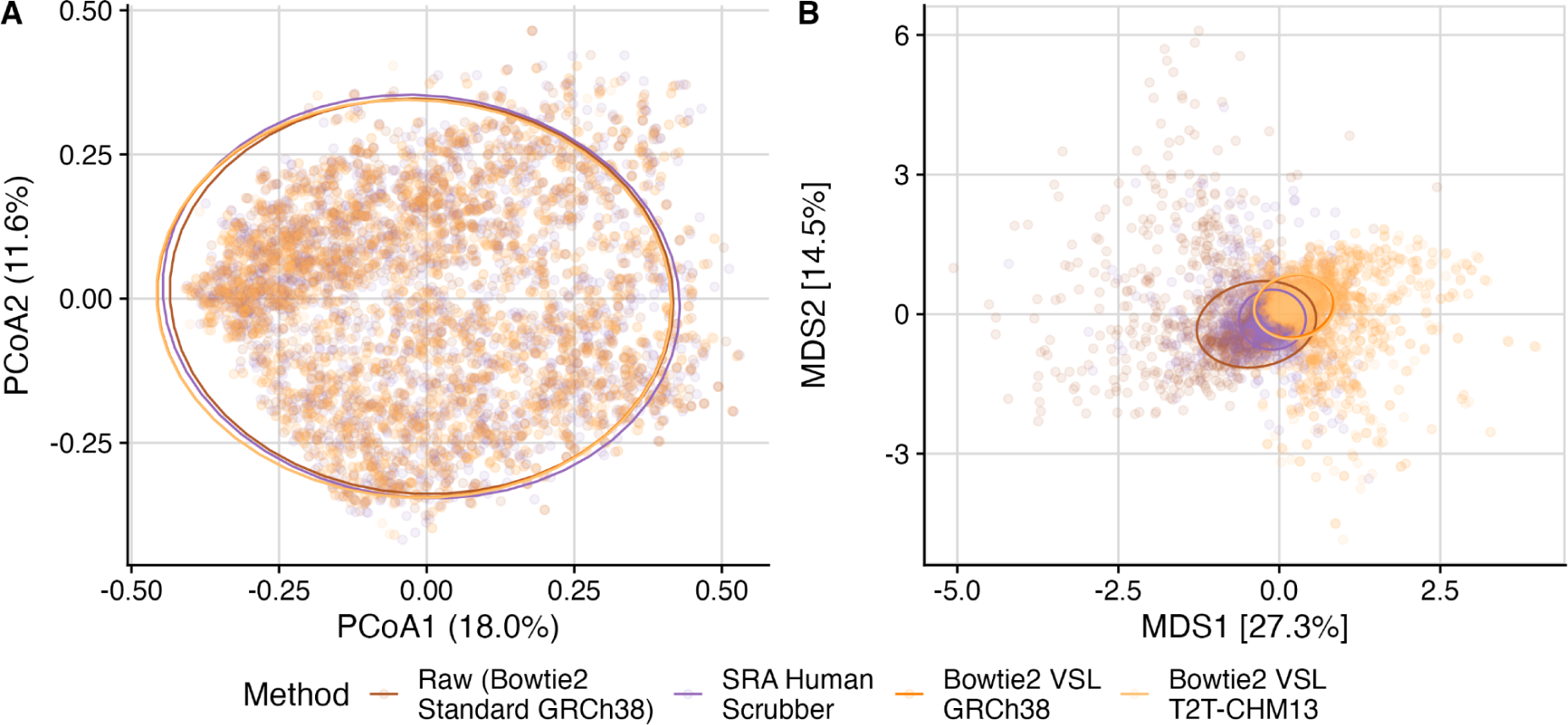
(A) a PCoA of the metagenomic data coloured by the different HRR methods. (B) an MDS visualising the differences in the data after removing the variation contributed by the sample.

**Fig. S4.**
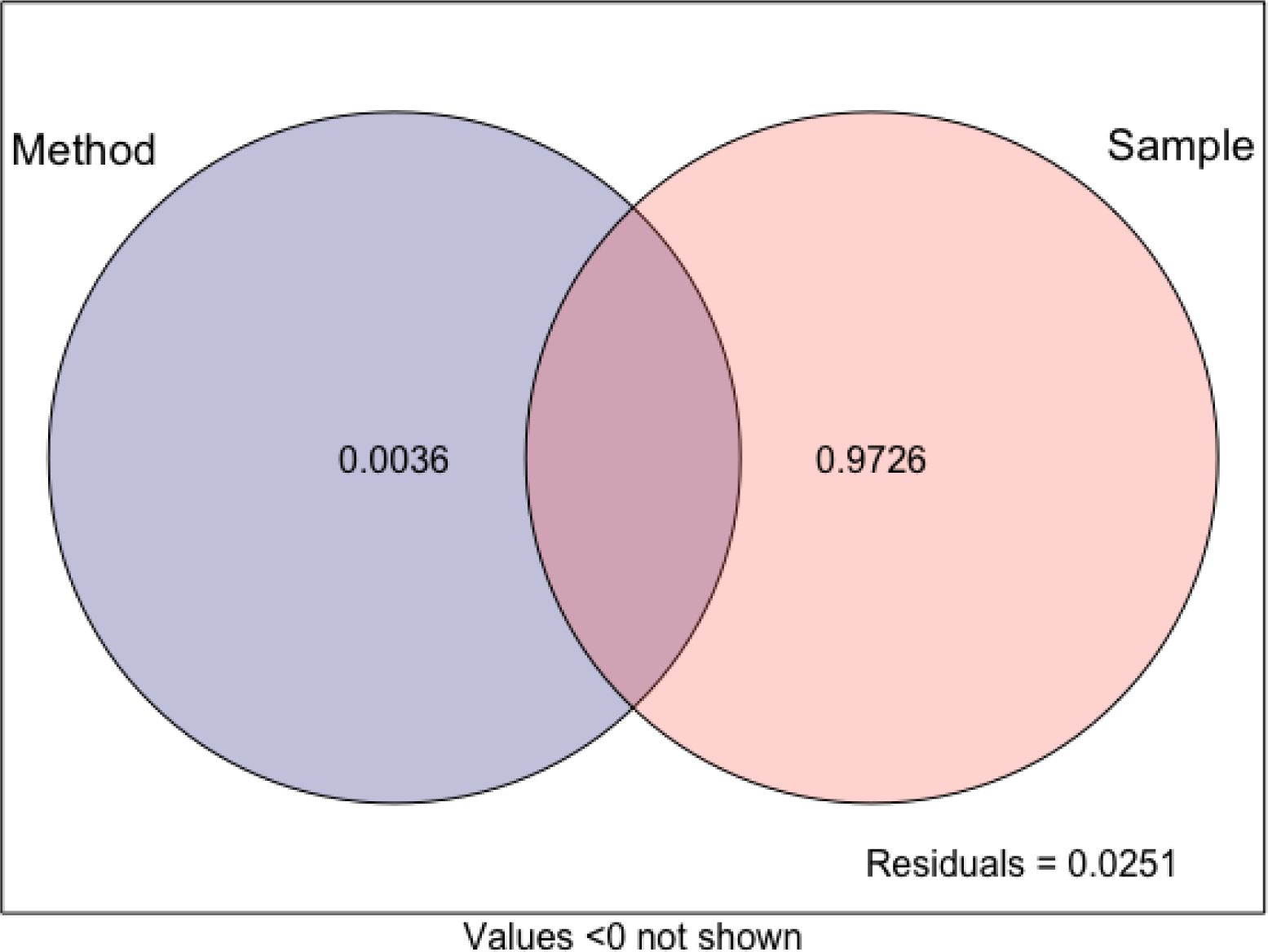
A Venn diagram visualising the variation partition within the metagenomic dataset.

**Fig. S5.**
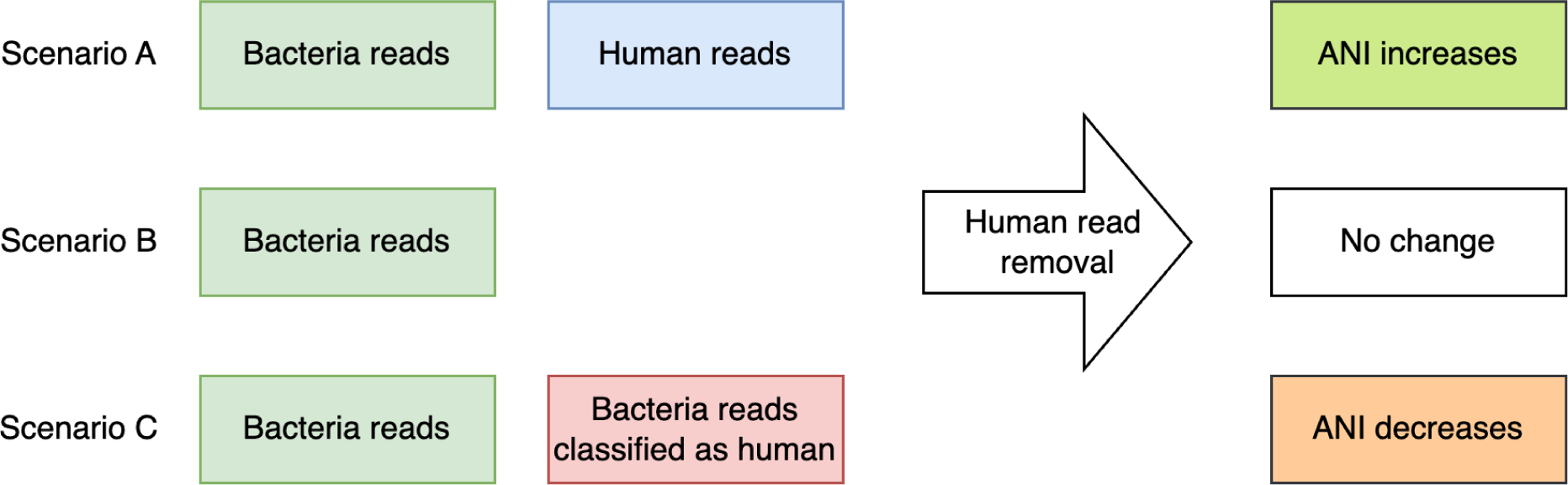
A flowchart illustrates how human read removal (HRR) affects average nucleotide identity (ANI) scores across different scenarios. ANI is calculated using Sylph, which utilises k-mers and is sensitive to changes in read composition. In Scenario A, ANI is calculated from both bacterial and human reads, where human reads resemble bacterial sequences, leading to a lower reported ANI score. Removing human reads increases the ANI score. In Scenario B, no human reads interfere with ANI calculation, so applying HRR has no effect. In Scenario C, some bacterial reads are misclassified as human reads and removed during HRR, reducing the total reads used for ANI calculation and lowering the ANI score.

**Table S1:**
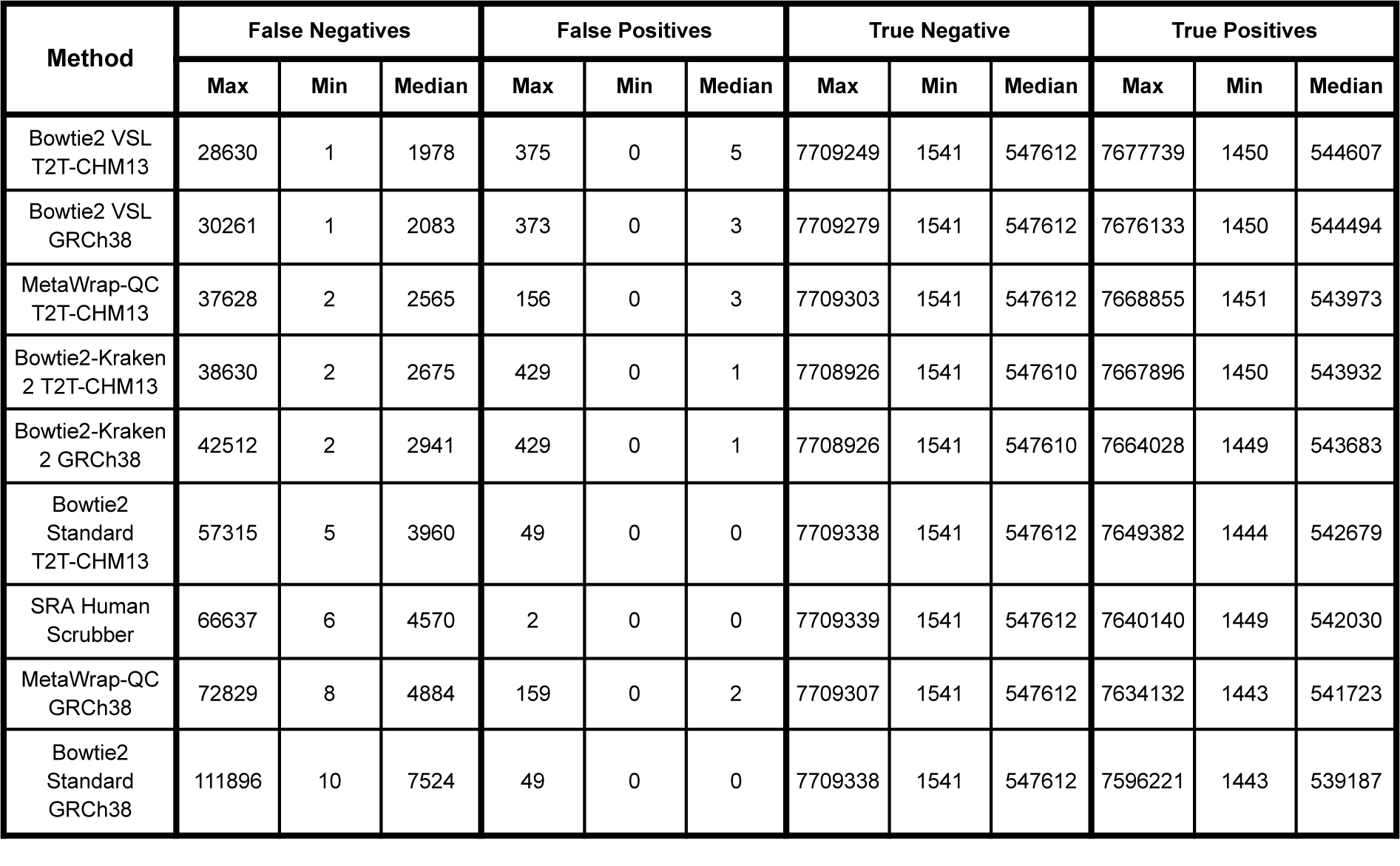
Maximum, minimum and median number of reads in each binary classification category across the synthetic mixture titration dataset. Rows are ordered by the Max False Negatives column.

**Table S2:** Median calculation for all the alpha diversity.

| Alpha diversity metric | Raw (Bowtie2 Standard GRCh38) | SRA Human Scrubber | Bowtie2 VSL GRCh38 | Bowtie2 VSL T2T-CHM13 |
| --- | --- | --- | --- | --- |
| Richness | 31 | 19 | 11 | 11 |
| Shannon | 2.03 | 1.89 | 1.84 | 1.84 |
| Inverse Simpson | 5.06 | 4.77 | 4.84 | 4.81 |
| Pielou's evenness | 0.62 | 0.68 | 0.78 | 0.78 |

**Table S3:** The delta mean of the beta coefficient obtained from a differential analysis.

| Phylum | Bowtie2 VSL<br>T2T-CHM13 vs<br>Bowtie2 VSL<br>GRCh38 | Bowtie2 VSL<br>T2T-CHM13 vs SRA<br>Human Scrubber | Bowtie2 VSL<br>T2T-CHM13 vs SRA<br>Human Scrubber |
| --- | --- | --- | --- |
| Actinomycetota | -0.009 | -0.192 | -0.182 |
| Bacillota | -0.002 | -0.202 | -0.199 |
| Bacillota_A | -0.013 | -0.167 | -0.154 |
| Bacteroidota | 0.007 | -1.021 | -1.027 |
| Pseudomonadota | 0.006 | -0.858 | -0.864 |
| Spirochaetota | 0.014 | -1.366 | -1.38 |

## References

1. Marotz CA, Sanders JG, Zuniga C, Zaramela LS, Knight R, Zengler K. Improving saliva shotgun metagenomics by chemical host DNA depletion. Microbiome. 2018 Feb 27;6(1):42.

2. Bush SJ, Connor TR, Peto TEA, Crook DW, Walker AS. Evaluation of methods for detecting human reads in microbial sequencing datasets. Microb Genom [Internet]. 2020 Jul;6(7). Available from: 10.1099/mgen.0.000393

3. Goig GA, Blanco S, Garcia-Basteiro AL, Comas I. Contaminant DNA in bacterial sequencing experiments is a major source of false genetic variability. BMC Biol. 2020 Mar 2;18(1):24.

4. Tomofuji Y, Sonehara K, Kishikawa T, Maeda Y, Ogawa K, Kawabata S, et al. Reconstruction of the personal information from human genome reads in gut metagenome sequencing data. Nat Microbiol. 2023 Jun;8(6):1079–94.

5. Lannelongue L, Grealey J, Inouye M. Green Algorithms: Quantifying the carbon footprint of computation. Adv Sci (Weinh). 2021 Jun 2;8(12):2100707.

6. Ulhuq FR, Barge M, Falconer K, Wild J, Fernandes G, Gallagher A, et al. Analysis of the ARTIC V4 and V4.1 SARS-CoV-2 primers and their impact on the detection of Omicron BA.1 and BA.2 lineage-defining mutations. Microb Genom. 2023 Apr 21;9(4):mgen000991.

7. Church DM, Schneider VA, Graves T, Auger K, Cunningham F, Bouk N, et al. Modernizing reference genome assemblies. PLoS Biol. 2011 Jul 5;9(7):e1001091.

8. Schneider VA, Graves-Lindsay T, Howe K, Bouk N, Chen HC, Kitts PA, et al. Evaluation of GRCh38 and de novo haploid genome assemblies demonstrates the enduring quality of the reference assembly. Genome Res. 2017 May;27(5):849–64.

9. Nurk S, Koren S, Rhie A, Rautiainen M, Bzikadze AV, Mikheenko A, et al. The complete sequence of a human genome. Science. 2022 Apr;376(6588):44–53.

10. The 1000 Genomes Project Consortium, Auton A, Abecasis GR, Altshuler DM (Co-Chair), Durbin RM (Co-Chair), Abecasis GR, et al. A global reference for human genetic variation. Nature. 2015 Oct 1;526(7571):68–74.

11. Langmead B, Salzberg SL. Fast gapped-read alignment with Bowtie 2. Nat Methods. 2012 Mar 4;9(4):357–9.

12. Constantinides B, Hunt M, Crook DW. Hostile: accurate decontamination of microbial host sequences. Bioinformatics [Internet]. 2023 Dec 1;39(12). Available from: 10.1093/bioinformatics/btad728

13. KneadData – The Huttenhower Lab [Internet]. [cited 2025 Feb 3]. Available from: https://huttenhower.sph.harvard.edu/kneaddata/

14. Wood DE, Lu J, Langmead B. Improved metagenomic analysis with Kraken 2. Genome Biol. 2019 Nov 28;20(1):257.

15. Uritskiy GV, DiRuggiero J, Taylor J. MetaWRAP-a flexible pipeline for genome-resolved metagenomic data analysis. Microbiome. 2018 Sep 15;6(1):158.

16. Katz KS, Shutov O, Lapoint R, Kimelman M, Brister JR, O’Sullivan C. STAT: a fast, scalable, MinHash-based k-mer tool to assess Sequence Read Archive next-generation sequence submissions. Genome Biol. 2021 Sep 20;22(1):270.

17. Di Tommaso P, Chatzou M, Floden EW, Barja PP, Palumbo E, Notredame C. Nextflow enables reproducible computational workflows. Nat Biotechnol. 2017 Apr 11;35(4):316–9.

18. Nicholls SM, Quick JC, Tang S, Loman NJ. Ultra-deep, long-read nanopore sequencing of mock microbial community standards. Gigascience [Internet]. 2019 May 1;8(5). Available from: 10.1093/gigascience/giz043

19. Brister JR, Ako-Adjei D, Bao Y, Blinkova O. NCBI viral genomes resource. Nucleic Acids Res. 2015 Jan;43(Database issue):D571–7.

20. Ju Y, Zhang Z, Liu M, Lin S, Sun Q, Song Z, et al. Integrated large-scale metagenome assembly and multi-kingdom network analyses identify sex differences in the human nasal microbiome. Genome Biol. 2024 Oct 8;25(1):257.

21. Shaw J, Yu YW. Rapid species-level metagenome profiling and containment estimation with sylph. Nat Biotechnol. 2024 Oct 8;1–12.

22. Parks DH, Chuvochina M, Rinke C, Mussig AJ, Chaumeil PA, Hugenholtz P. GTDB: an ongoing census of bacterial and archaeal diversity through a phylogenetically consistent, rank normalized and complete genome-based taxonomy. Nucleic Acids Res. 2022 Jan 7;50(D1):D785–94.

23. R Core Team. R: A Language and Environment for Statistical Computing [Internet]. Vienna, Austria: R Foundation for Statistical Computing; 2021. Available from: https://www.R-project.org/

24. Oksanen J, Simpson GL, Blanchet FG, Kindt R, Legendre P, Minchin PR, et al. https://vegandevs.github.io/vegan/. 2025. *vegan: Community Ecology Package* (R package version 2.7-0).

25. Lahti L, Shetty S. Bioconductor. 2017 [cited 2025 Mar 7]. microbiome. Available from: http://bioconductor.org/packages/microbiome/

26. Arbizu PM. pairwiseAdonis: Pairwise multilevel comparison using adonis [Internet]. Github; 2020 [cited 2025 Mar 7]. Available from: https://github.com/pmartinezarbizu/pairwiseAdonis

27. Brunson C. corybrunson/ggalluvial: faceting bug fix [Internet]. Zenodo; 2020 [cited 2025 Mar 14]. Available from: https://zenodo.org/records/4012484

28. Mallick H, Rahnavard A, McIver LJ, Ma S, Zhang Y, Nguyen LH, et al. Multivariable association discovery in population-scale meta-omics studies. PLoS Comput Biol. 2021 Nov 16;17(11):e1009442.

29. Paradis E, Schliep K. ape 5.0: an environment for modern phylogenetics and evolutionary analyses in R. Bioinformatics. 2019 Feb 1;35(3):526–8.

30. Aganezov S, Yan SM, Soto DC, Kirsche M, Zarate S, Avdeyev P, et al. A complete reference genome improves analysis of human genetic variation. Science. 2022 Apr 1;376(6588):eabl3533.

31. Rhie A, Nurk S, Cechova M, Hoyt SJ, Taylor DJ, Altemose N, et al. The complete sequence of a human Y chromosome. Nature. 2023 Sep 23;621(7978):344–54.

32. Kim D, Paggi JM, Park C, Bennett C, Salzberg SL. Graph-based genome alignment and genotyping with HISAT2 and HISAT-genotype. Nat Biotechnol. 2019 Aug 2;37(8):907–15.

